# A Scalable Optimization Algorithm for Solving the Beltway and Turnpike Problems with Uncertain Measurements

**DOI:** 10.1101/2024.02.15.580520

**Authors:** C.S. Elder, Minh Hoang, Mohsen Ferdosi, Carl Kingsford

## Abstract

The Beltway and Turnpike problems entail the reconstruction of circular and linear one-dimensional point sets from unordered pairwise distances. These problems arise in computational biology when the measurements provide distances but do not associate those distances with the entities that gave rise to them. Such applications include molecular structure determination, genomic sequencing, tandem mass spectrometry, and molecular error-correcting codes (since sequencing and mass spec technologies can give lengths or weights, usually without connecting them to endpoints). Practical algorithms for Turnpike are known when the distance measurements are accurate, but both problems become strongly NP-hard under any level of measurement uncertainty. This is problematic since all known applications experience some degree of uncertainty from uncontrollable factors. Traditional algorithms cope with this complexity by exploring a much larger solution space, leading to exponential blowup in terms of both time and space. To alleviate both issues, we propose a novel alternating optimization algorithm that can scale to large, uncertain distance sets with as many as 100,000 points. This algorithm is space and time-efficient, with each step running in *O*(*m* log(*m*)) time and requiring only 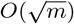 working space for a distance set of size *m*. Evaluations of this approach on synthetic and partial digest data showcase improved accuracy and scalability in the presence of uncertain, duplicated, and missing distances. Our implementation of the algorithm is available at https://github.com/Kingsford-Group/turnpikesolvermm.

## 1. Introduction

The Turnpike problem is to reconstruct *n* unknown points on a line from all 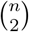 pairwise distances between them provided without labels. The Beltway problem is a variant where the points lie on a circle instead of a line. These problems arise frequently in computational biology applications when the measurement (e.g. mass spectrometry or sequencing) provides distances but does not associate those distances with the entities that gave rise to them. In the tandem mass spec application, for example, when sequencing an unknown peptide *a*_1_*a*_2_ … *a*_*n*_, the “points” are the weights of the fragments *a*_1_, *a*_1_*a*_2_, *a*_1_*a*_2_*a*_3_, … — reconstructing those points would suggest the identity of each amino acid *a*_*i*_ via its weight. However, the MS/MS measurement provides the weights of every fragment *a*_*i*_ … *a*_*j*_ (which can be treated as “distances” between points *i* and *j*) without associating that measurement with *i* or *j*. Various heuristics for this problem are applied to structure estimation of biomolecules [12], *de novo* sequencing of linear and cyclic peptides [17,8], and reconstructing DNA sequences from their partially digested fragments [23,21]. Some versions of the problem provide additional labeling information and reduced distance subsets, such as the labeled partial digest problem that separates the endpoint distances from the all-pairs distance set and the simplified partial digest problem [4] that returns only a subset of the distances. Other variants of the Turnpike problem are applied to quantum phase estimation [27] and molecular error-correcting codes used for databases [10].

The Exact Turnpike variant of the problem, where all distances are observed without error, can be solved exactly via a backtracking algorithm that alternates between placing the largest remaining distance and matching derived and ground truth distances [22]. While there exist pathological cases for which it incurs exponential runtimes [26], this algorithm is generally efficient in practice with an expected run time of *𝒪* (*n*^2^ log *n*) for random instances [21]. Various extensions to the backtracking algorithm have been proposed to improve both expected and worst case runtimes, including a variant based on breadth-first search [1], that are empirically faster. One example efficiently solves known pathological instances [18]. In contrast, algorithms for Beltway are not efficient, with a worst case runtime of *𝒪* (*n*^*n*^ log *n*) that is often realized in practice [8]. Other approaches to Turnpike include a fixed-parameter tractable algorithm that works by factoring a polynomial and scales with the largest distance in the set [13]. Regrettably, this approach is highly susceptible to numerical precision errors [14]. This restricts its practical applicability whenever floating point arithmetic is used. A semidefinite relaxation also exists that is able to solve some instances but is known to be numerically unstable and suffer from runtimes far exceeding the backtracking algorithm in practice [12].

When the distance measurements are uncertain, we have the Noisy Turnpike and Noisy Beltway problems, which are both strongly NP-complete as demonstrated by a reduction from the three-partition problem [6]. Skiena et al. modified the backtracking approach to use intervals instead of points to accommodate measurement uncertainty [21], but these modifications lead to the consideration of exponentially many paths, limiting the algorithm’s efficiency and practical applicability [12]. Pandurangan et al. assumed that the partial digestion results from both ends of the double stranded DNA sample are observed [20]. Fomin et al. performed the equivalent modifications for Noisy Beltway and reduce the running time by removing redundant measurements, but as with the exact case, only very small Noisy Beltway instances can be solved with this algorithm [9]. More recently, Huang et al. model both the Noisy Turnpike and Noisy Beltway problems as probabilistic inference of the point assignments using discrete bins that quantize the input domain [12]. In this approach, the bin size is set to be smaller than the smallest distance, and hence it was assumed that no bin can contain more than one point. However, this only holds true when the observation error is sufficiently small in magnitude relative to the smallest distances. As such, the accuracy of this algorithm deteriorates in noisier instances. In addition, it also struggles to efficiently solve larger problem instances, with *n* > 500 out of reach at present.

In section 2, we propose a novel approach to solving the Noisy Turnpike and Noisy Beltway problems using a bilevel optimization scheme that alternates between estimating the point-distance correspondence and recovering the original point set given this assignment. Our formulation’s non-convex optimization landscape contains many saddle points and local optima. We accommodate for this by introducing a divide-and-conquer step to recursively correct small-scale mistakes that lead to low-quality solutions. Our algorithm runs in time *𝒪* (*n*^2^ log *n*) for each step, with time dominated by a low-cost sorting step.

In Section 3, we empirically demonstrate the performance of our proposed Minorization-Maximization algorithm (MM) in various synthetic and realistic biological settings, such as the partial digestion task [12]. Our algorithm arrives at highly accurate solutions even in extremely Noisy observation conditions. We also demonstrate that the proposed algorithm runs more efficiently than previous approaches and empirically matches our theoretical runtime expectation. Most notably, the proposed algorithm can efficiently process partial digestion instances with up to a hundred thousand digested fragments, which is realistically on the scale of a whole genome and has never been achieved by previous methods. Moreover, we provide an extension of the method in Appendix C to problem variants that provide additional labeling information and reduced distance sets. In summary, this algorithm advances the capacity to address both the Noisy Turnpike and Noisy Beltway problems, and thereby improves the accuracy and scalability of various biological tasks that make use of these formulations.

## 2 Method

### 2.1 Problem Setting

Let *m* = *n*(*n* − 1)/2 and *D* ∈ ℝ^*m*^ be a vector of pairwise distances between *n* points. We denote the ground truth vector containing the points to be recovered as *z* ∈ ℝ^*n*^. Without loss of generality, we assume that 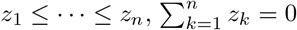, and ||*z*||_2_ = 1; that is, the unknown points are named in sorted order, centered around zero, and have unit norm. These assumptions do not fundamentally change the problem, but are nonetheless important as they prevent trivial non-uniqueness. The first and second assumptions hold because the distance set is invariant to translation and permutation of the points, allowing us to look for a centered, sorted solution vector *z*. The third assumption follows because we can construct a scaled distance set 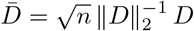 from the original distances that generates 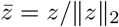.

We are able to perform this scaling because

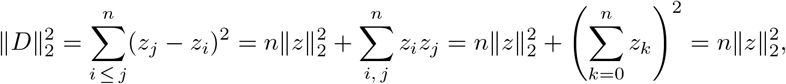

where we used the fact that *z* is centered. Let *Ƶ* be the set of vectors in ℝ ^*n*^ that satisfy all three assumptions above (treating dimensions of vectors in *Ƶ* as the point locations), and let *𝒮*_*m*_ denote the set of all permutation matrices of *m* items. We use Turnpike to refer to both the Exact Turnpike and Noisy Turnpike variants, when statements apply to both. The Exact Turnpike problem is formalized as finding a vector 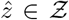 such that 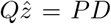 for some *P* ∈ *𝒮*_*m*_, and where *Q* ∈ ℝ^*m×n*^ is a fixed incidence matrix defined as follows. Each row in *Q* corresponds to a pair of indices *j* > *i*. For convenience, we let the function *α*(*i, j*) map the index pair (*i, j*) to its (arbitrary) row index in *Q*. The incidence matrix *Q* is constructed such that *Q*_*α*(*i,j*),*j*_ = 1 and *Q*_*α*(*i,j*),*i*_ = −1 are the only non-zero entries in *Q*_*α*(*i,j*)_. It follows that 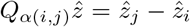, and 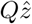 contains all the pairwise distances generated by 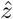. Furthermore, if 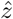 recovers the ground truth *z*, then 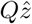 must also be a permutation of *D*, which explains the role of *P* in the objective above. In the Noisy Turnpike case, which is the focus of this paper, exact recovery is not possible in general due to the corrupted observations. Therefore, the objective can be written in the form of an optimization task:

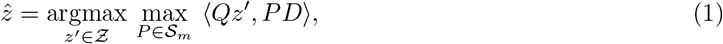

where ⟨·, ·⟩ denotes the inner product of two vectors. This is equivalent to minimizing the *𝓁*_2_ distance between *Qz*^*′*^ and *PD* because the norm of *P, Q* and *D* are constant, *z*^*′*^ is normalized based on our previous assumptions, and optimality in the exact case will take place when *Qz*^*′*^ = *PD*. Nonetheless, Eq. (1) is more convenient for our subsequent derivation.

### 2.2 Minorization-Maximization scheme for optimizing the Turnpike objective

#### Algorithm 1

Minorization-Maximization (MM)

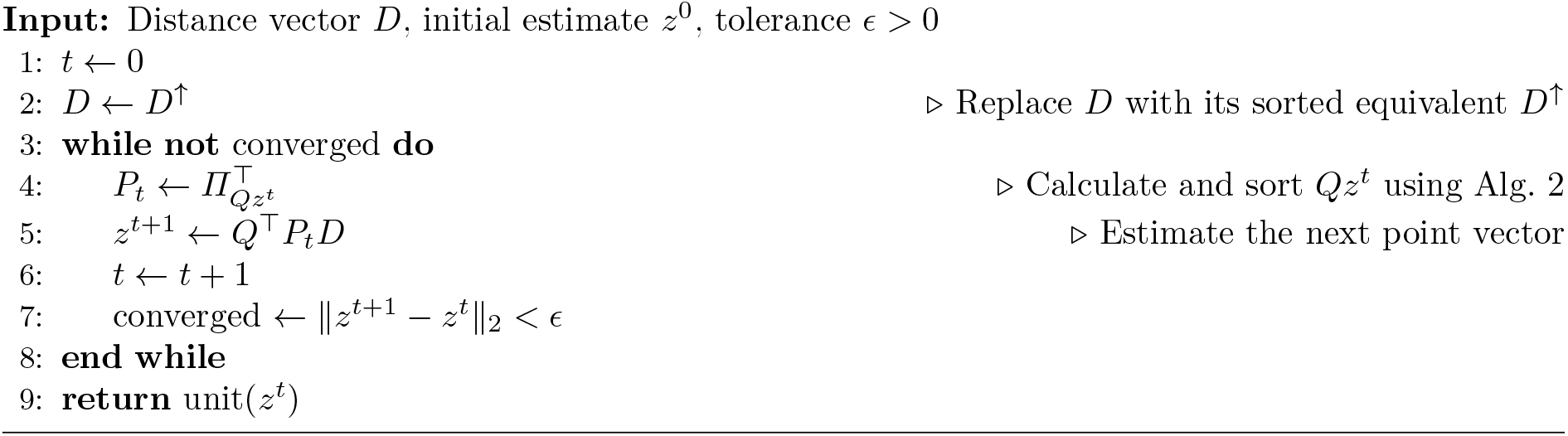

We first observe that Eq. (1) is bilinear because fixing either *P* or *z* reduces the problem to a linear function. This motivates a bilevel minorization-maximization (MM) scheme [24] to optimize this objective. In particular, we relax our objective into two alternating subproblems. At iteration *t* + 1, for *t* > 0, the first subproblem fixes an estimation *z*^*t*^ and solves for

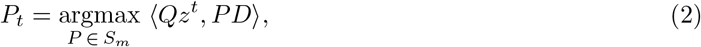

which has a closed-form solution, as shown in Proposition 1 below. This closed form is a variant of the rearrangement inequality [11], a connection used in Lemma 1.

#### Lemma 1.

*Suppose y* = ⟨*y*_1_, *y*_2_, …, *y*_*n*_⟩ ∈ ℝ^*n*^ *is a sorted vector, i*.*e*., *y*_1_ ≤ *y*_2_ ≤ … ≤ *y*_*n*_. *The problem*

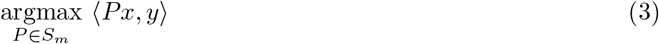

*is maximized by any permutation P*_*x*_ *such that P*_*x*_*x is in sorted order*.

*Proof*. Let *P* ∈ *𝒮*_*m*_ be a non-sorting permutation, meaning there exist indices *i* < *j* where (*Px*)_*i*_ > (*Px*)_*j*_. Notice that transposing elements *i* and *j* increases the objective because

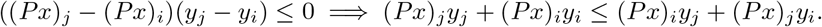

Thus the permutation that first applies *P* then transposes elements *i* and *j* is no worse than *P*. Iterating this argument leads to a sorting permutation that is also no worse than *P*. As the initial permutation was arbitrary, this shows that there exists a globally maximizing permutation that sorts *x*. To finish the proof, notice that any two sorting permutations *P*_1_ and *P*_2_ must have the same objective value since sortedness implies 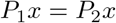.

#### Proposition 1.

*Let Π*^⊤^ *be a permutation that puts Qz into sorted order. The permutation Π is a globally maximizing solution to Eq*. (2).

*Proof*. Without loss of generality, we assume that *D* is sorted as a preprocessing step to the Noisy Turnpike problem. We can rewrite the expression in Eq. (2) as ⟨*Qz*^*t*^, *PD*⟩ = ⟨*P*^⊤^(*Qz*^*t*^), *D*⟩. By Lemma 1, any permutation *Π*^⊤^ that sorts *Qz* must be a global maximizer of the right-hand side. Notice the transpose *Π* is the equivalent solution on the left-hand side, proving the claim.

On the other hand, the second subproblem fixes an estimation for *P*_*t*_, which is the closed-form solution derived above, and solves for:

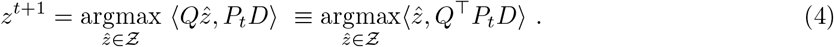

Since the inner product of two vectors is maximized when they are parallel and 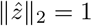 by assumption, the maximum objective value is obtained when 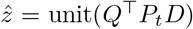, where unit(·) scales a vector to unit norm.

As objectives (2) and (4) have closed-form solutions, they motivate a practical bilevel optimization routine described in Alg. 1. Note that the unit projection does not affect the permutation in the next iteration, so we omit it until the vector is returned. We avoid storing both the incidence matrix and intermediate distance vector by using implicit matrix multiplication and a problem-specific matching algorithm Alg. 2. The runtime of the optimization inner loop is derived in Proposition 1. In the same proposition, we also derive a memory efficient implementation that avoids storing intermediate values during optimization.

#### Lemma 2.

*The priority queue in Alg. 2 uses z*^*t*^ *interval order, i*.*e*., (*i, j*) ≤ (*i*^*′*^, *j*^*′*^) ⟺ (*z*^*t*^[*j*] − *z*^*t*^[*i*]) ≤ (*z*^*t*^[*j*^*′*^] − *z*^*t*^[*i*^*′*^]). *This ordering satisfies i* ≤ *i*^*′*^, *j*^*′*^≤ *j and implies* (*i, j*) ≤ (*i*^*′*^, *j*^*′*^) *when z*^*t*^ *is sorted*.

*Proof*. For *i* ≤ *i*^*′*^ and *j*^*′*^ ≤ *j*, we have from sortedness that *z*^*t*^[*i*] ≤ *z*^*t*^[*i*^*′*^] and *z*^*t*^[*j*^*′*^] ≤ *z*^*t*^[*j*], which implies *z*^*t*^[*j*^*′*^] − *z*^*t*^[*i*^*′*^] ≤ *z*^*t*^[*j*] − *z*^*t*^[*i*]. This is the definition of (*i*^*′*^, *j*^*′*^) ≤ (*i, j*).

#### Proposition 2.

*Upon termination of Alg. 2, the vector z*^*t*+1^ *contains* 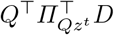. *Moreover, the algorithm runs in O*(*n*^2^ log *n*) *time and uses only 𝒪* (*n*) *nonnegative integers for non-constant storage, where n is the number of points*.

*Proof*. We first prove that the priority queue pops the *t*^th^ smallest distance during iteration *t*. To that end, we define the sequences 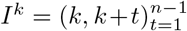. We note that the sequences *I*^1^, …, *I*^*n*−1^ partition the 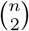 possible intervals. Lemma 2 establishes that these chains are in interval sorted order, and thus implies Alg. 2 produces the smallest unseen interval at each iteration. This holds because the queue holds the smallest element from each sequence, and we add the next one until each sequence has been exhausted.

By the discussion above, during iteration *t*, (*i, j*) is the (potentially non-unique) *t*^th^ smallest interval. Thus the sorting permutation will send interval (*i, j*) to index *t*, and its transpose (i.e., inverse) will send index *t* to interval (*i, j*), implying *D*[*t*] will be used as the (*i, j*) distance. When multiplying by *Q*^⊤^, the (*i, j*) distance entry contributes only to the *i*^th^ and *j*^th^ points. Specifically, the (*i, j*) entry is subtracted from *z*^*t*+1^[*i*] and added to *z*^*t*+1^[*j*], which is immediately performed in Alg. 2’s loop. Thus the algorithm terminates with *z*^*t*+1^ = *Q*^⊤^*P*_*t*_*D* since we initially zero it out and accumulate all the entries that contribute to it.

The algorithm performs *𝒪* (*n* log *n*) work before the main loop. The main loop performs *𝒪* (*n*^2^ log *n*) work because of the priority queue, which takes *𝒪* (log *n*) time per iteration using standard implementations. This is unaffected by the interval comparison function, as that performs only constant work. The priority queue uses the only non-constant memory, as it needs to store *𝒪* (*n*) non-negative integers for the intervals.

#### Proposition 3.

*The outer loop of Alg. 1 must terminate in a finite number of steps. The inner-loop runs in time 𝒪* (*n*^2^ log *n*) *and requires only 𝒪* (*n*) *non-constant storage*.

*Proof*. For the first claim, notice that the sorting permutation in each iteration fully decides the point vector that is produced in the next step, meaning the set of possible output vectors is finite. From our earlier derivation, we also know that the permutation and point vector must improve the objective at each step. This means the algorithm cannot continue to make progress indefinitely and will terminate when *z*^*t*^ = *z*^*t*+1^.

For the second claim, notice that steps 4 and 5 of Alg. 1 take *O*(*n*^2^ log *n*) time and need only *𝒪* (*n*) non-negative integers of non-constant storage by the result of Proposition 1. We also only need to keep two *n* floating point vectors to calculate step 7. □

In practice, we observed that Alg. 1 converges quickly but is prone to becoming trapped in local maxima. To prevent this, we further propose a divide-and-conquer heuristic formally described in Alg. 3. In particular, after each pass of Alg. 1, we partition the estimation 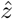 into non-overlapping subsets 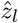 and 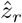. In our implementation, the median is used to form the partitions, but any rule works with this framework. No matter the choice, this segments the distance set *D* into three portions: (a) *D*_*ll*_ contains the distances among points in 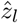; (b) *D*_*rr*_ contains the distances among points in 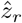; and (c) *D*_*lr*_ contains the distances between pairs of points respectively in 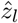 and 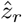. Even though we do not have the ground truth assignment of the point-distance correspondence, we can use the estimated permutation matrix *P* to perform this segmentation.

The intuition behind our proposed heuristic is that, if 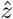 and *P* are the optimal Turnpike solution, then subsequent applications of Alg. 1 on (*z*_*l*_, *D*_*ll*_) and (*z*_*r*_, *D*_*rr*_) will not alter this solution. Otherwise, the recursive sub-routines will likely not get trapped in the same local maxima as the parent routine and will serve as a self-correcting mechanism for Alg. 1 by returning the adjusted permutations *P*_*l*_ and *P*_*r*_. At this point, we can adjust 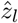 and 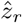 by solving the following regression tasks:

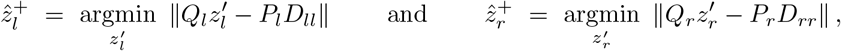

where *Q*_*l*_ and *Q*_*r*_ are the respective incidence submatrices corresponding to 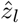 and 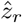. To avoid storing the incidence matrices, we can use any matrix-free solver such as the conjugate gradient method [25] (see Alg. 4 in Appendix A for the matrix-free oracles). As the adjusted estimation 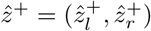 breaks away from potential local maxima, this routine is repeated until convergence, as described in Alg. 3 below. We provide a visualization of how this improves solutions in Fig. 3 (see Appendix B).

#### Algorithm 2

*Q*^⊤^ applied to matched *D*

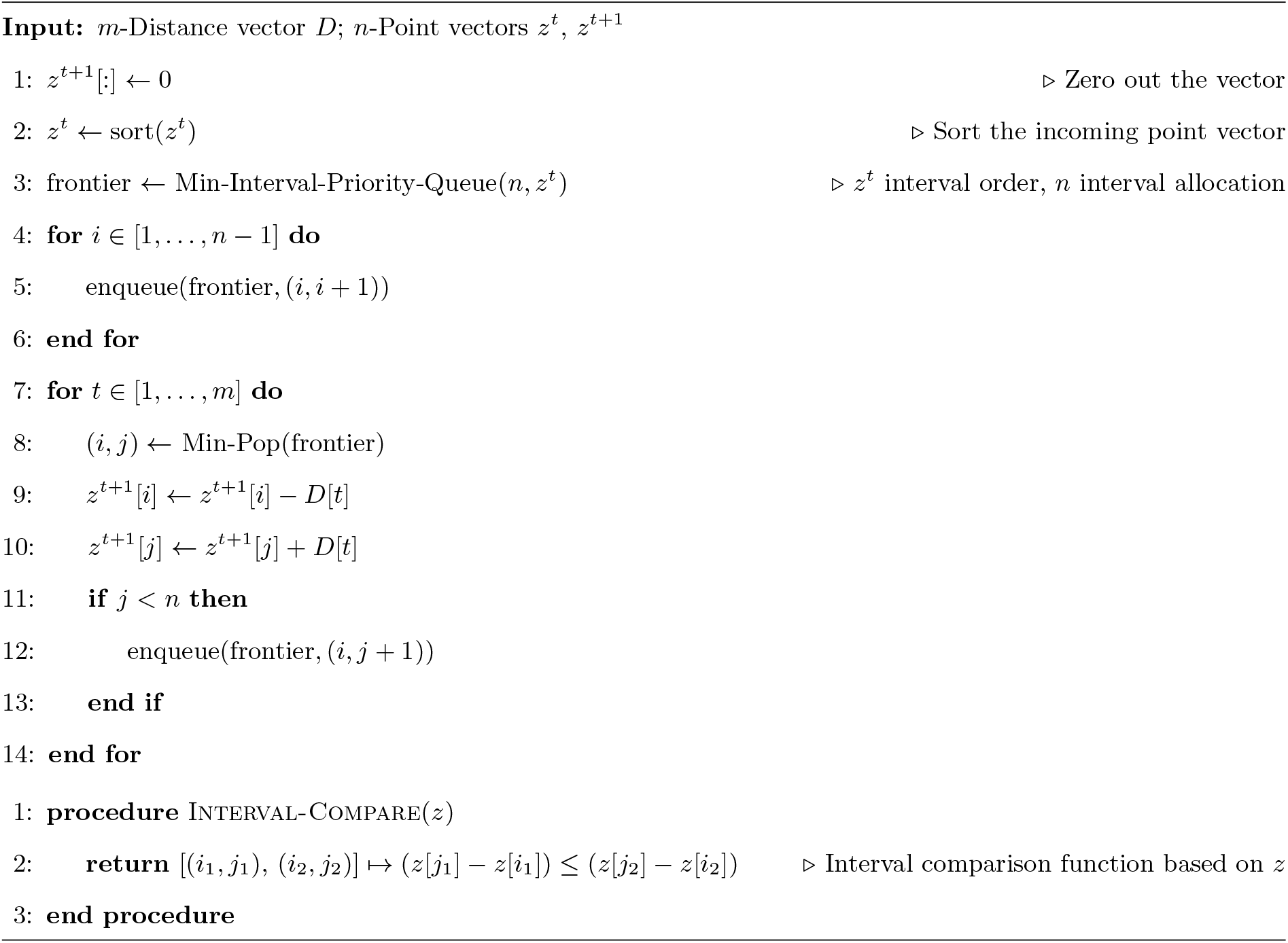

#### Remark

An alternative approach (which will be compared against our method in Section 3) to solving Eq. (1) applies Birkhoff’s theorem [3], which states that the polytope *ℬ*_*m*_ of *m* × *m* doubly stochastic matrices is the convex hull of *𝒮*_*m*_. This motivates a relaxation of Eq. (1) to optimize for *P* on *ℬ*_*m*_:

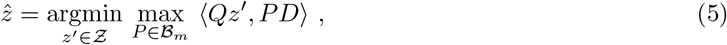

allowing for a differentiable permutation learning framework that combines (a) stochastic gradient descent over the space of square matrices; and (b) projection onto *ℬ*_*m*_ with the Sinkhorn operator [16]. In the case of the Turnpike and Noisy Turnpike problems, this approach requires the algorithm to optimize an *m*×*m* matrix, which holds an infeasibly large *Θ*(*n*^4^) entries. We refer to this alternative as the “gradient descent” method in the results below.

##### Algorithm 3

Minorization-Maximization Divide-and-Conquer (MMDQ)

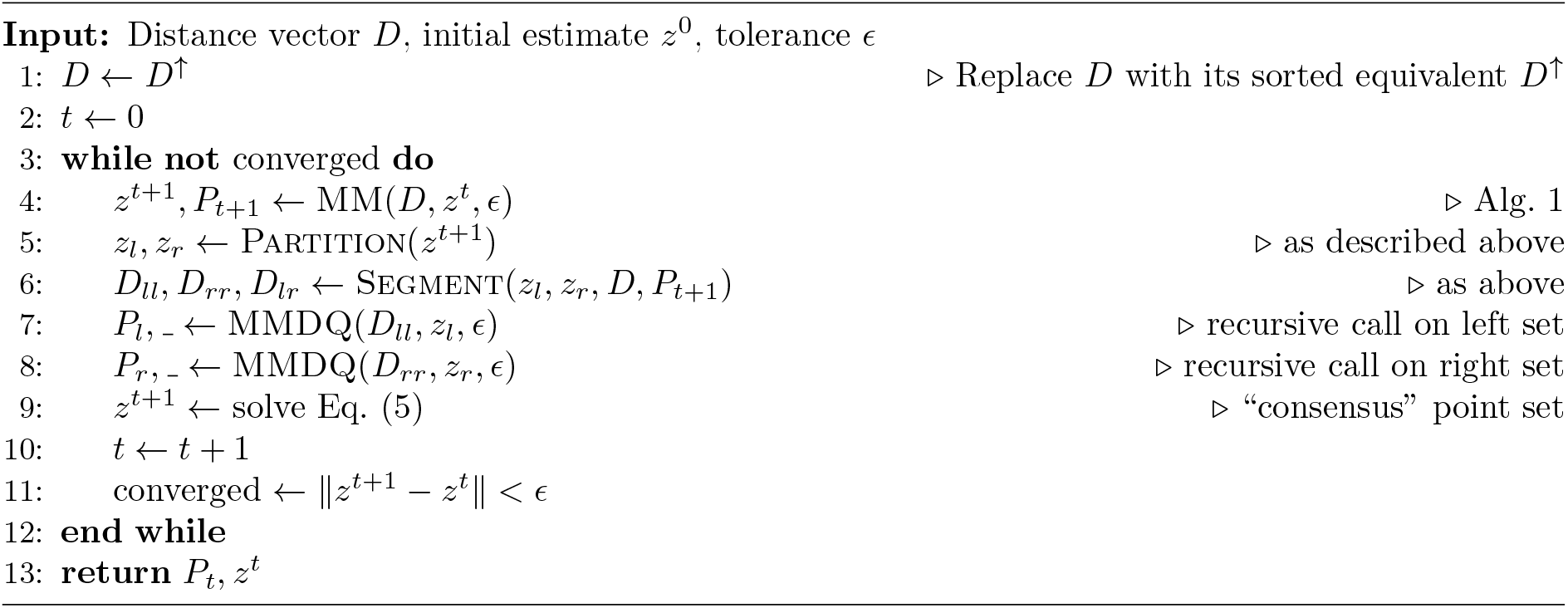

### 2.3 Extension to other variants

In the Beltway problem, we are given *n*(*n* − 1) unlabeled arc lengths (distances) between *n* points *p*_1_, …, *p*_*n*_ on a circle. Note that we receive double the number of distances as in the Turnpike case because there are two different arcs between any two points (i.e., clockwise and counter-clockwise). We only consider problem instances on a unit circle as we are able to scale the points using the transformation detailed in Huang et al. [12]. The Beltway problem can be solved within our framework (Section 2.2) with minor modifications. We also extend our method to a general variant of the Turnpike problem that handles both labeled and missing distances. This extension captures the labeled partial digest problem [20] and simplified partial digest problem [4]. See Appendix C for details.

### 2.4 Initializer Sampling

The choice of an initializer for Alg. 3 plays a critical role in achieving good convergence and overall performance. A well-chosen initializer can lead to faster convergence, improved stability, and a more accurate solution. Here, we consider three practical initializing schemes. The first scheme samples a random Gaussian vector and sorts it. Though efficient to implement, this scheme is unlikely to produce a good initializer if the ground set exhibits pathological features such as having spread-out point clusters. On the other hand, if the points are well-spread, this scheme often finds a close starting point. The second scheme provides a random permutation *P*_0_ to the sub-problem in Eq. (2) and sets *z*^0^ as its closed-form solution. This incorporates the combinatorial nature of Turnpike and potentially encourages more diverse exploration of the solution space. Nevertheless, selecting random permutations does not guarantee proximity to the optimal solution or even proximity to a valid distance permutation. The final scheme is a greedy-search method inspired by the classical backtracking approach [21]. That is, we sequentially fit the largest distance in *D* onto a line segment configuration (i.e., placing a new point to the left or to the right end of the segment based on this distance). However, unlike the original formulation—which uses backtracking to find the optimal placement—we make greedy choices to generate an initializer that will be polished with our algorithm afterwards, thus avoiding potentially exponential runtime.

## 3 Empirical results

### Experimental design

We assessed the performance of our proposed algorithm on the Turnpike, Beltway, and Labeled Partial Digest problems. As a baseline, we used synthetic data to validate our proposed algorithm’s performance on uncertain measurements and compared it to the backtracking method [21], the distribution matching method [12], and our projected gradient descent baseline using the Gumbel-Sinkhorn relaxation [16]. To evaluate the performance of our method in genome reconstruction, we conducted a series of experiments that simulated the reconstruction of a DNA sequence from fragments generated by enzymes. All experiments were implemented in Python 3.10 using a C++20 library implementing the algorithm integrated with Python using PyBind11 and conducted on a computer equipped with 1.0 TB of RAM, two Intel Xeon E5-2699A v4 CPUs, and a GTX 3080 GPU.

### Synthetic data

Synthetic datasets were generated by sampling *n* points on the real line from three distributions: the Cauchy distribution, the standard normal distribution, and the uniform distribution on [0, 1]. The uniform distribution was chosen to align with the setting explored by Huang et al. [12]. The normal distribution was chosen to test point sets with clustered points. The Cauchy distribution was selected to test the effect of contrasting scales and dispersion of points sampled from the Cauchy distribution present a challenge for *𝓁*_2_ optimization methods, which struggle with outlying values [5].

We examined sample sizes ranging from 50 to 2000 points (in increments of 50) and three additional large sample sizes of 5000, 10,000, and 100,000 to demonstrate the method’s scalability. To simulate measurement uncertainty of magnitude *ϵ* = 10^−*k*^ for integer *k* ∈ [1, 12], we added a Gaussian noise vector *g* ∼ *𝒩* (**0**, *ϵ***I**) to the given vector of pairwise distances [7]. We rounded the distance to zero when the amount of uncertainty exceeded the magnitude of the distance, which simulates missing distances. We predicted the points for each set of distribution, size, and uncertainty for 10 independent test cases. We run each algorithm 10 times and output the best estimate, which we quantified with the *𝓁*_2_ distance between the estimated and uncertain distance sets (the algorithms are deterministic, but the choice of initializer is random as described above). We recorded the mean absolute error (MAE) and mean squared error (MSE) between the estimated and ground point sets. The MAE is a continuous alternative to the binning distance [12] and is more suitable for our method since we do not explicitly assign points to bins.

Since the distribution matching method produces bins as its output, we use the midpoint of each bin as the predicted point.

### Study of different initialization schemes

We investigated the three initialization strategies (Section 2.4) to select one for subsequent experiments. We boosted the Gaussian point vector and permutation point vector initializer by drawing n distinct samples that were scored by solving Problem 2 for each and taking the maximum value. The sample with the maximum score from each strategy was used as the initializer. We used the Gaussian initializer as the starting point for the greedy-search initializer. We tested the strategies across all settings described previously. Fig. 1 shows the cosine distances between the estimated and uncertain distance vectors. Among the three approaches, the permutation strategy exhibited the worst similarity scores, with an average magnitude 13 times larger than that of the greedy-search strategy. The Gaussian strategy demonstrated an average error magnitude that was 8 times larger than the greedy-search initialization.

**Fig. 1:**
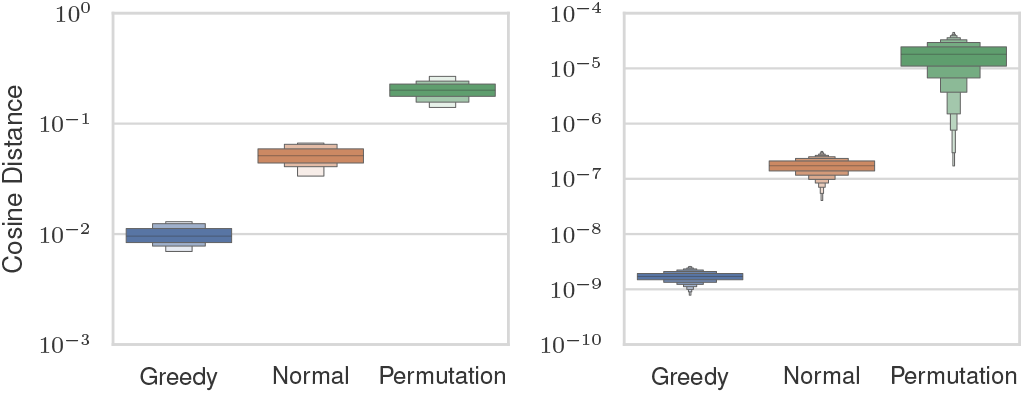
Cosine similarities between estimated and ground distance vectors (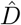 and *D*) before (left) and after (right) MM optimization for three different initialization schemes.

A better initial score does not necessarily guarantee a better reconstruction after optimization. To assess the efficacy of each initializer, we analyzed whether lower pre-optimization errors translated to reduced post-optimization errors. The cosine distance after optimization is also shown in Fig. 1. The permutation initialization had the highest errors and the greedy-search approach had the lowest errors, which is consistent with the pre-optimization cosine distance. We used the greedy-search initializer for our experiments since it exhibited the lowest post-optimization distance. The greedy-search strategy’s lower error comes at a computational cost. Table 1 shows the median runtime for the greedy-search initializer, Gaussian initializer, and optimization loop across a representative set of problem sizes. For all sizes, the greedy-search initialization takes more time than running the optimization, whereas the Gaussian initialization strategy runs in an order of magnitude less time than the optimization. This is due to the inherently serial nature of the greedy-search initializer, which requires all previous steps to be considered first. This is in contrast to the optimization loop, which has a runtime dominated by sorting, which is parallelized.

**Table 1:** Median runtimes (in seconds) for the MM optimizer, Gaussian and Greedy initializations over different sample sizes.

| Points | 100 | 500 | 1000 | 1500 | 2000 |
| --- | --- | --- | --- | --- | --- |
| Optimizer | 0.40 | 8.40 | 16.49 | 25.39 | 42.39 |
| Gaussian | 0.54 | 2.54 | 4.63 | 7.53 | 18.53 |
| Greedy | 0.23 | 17.56 | 36.49 | 55.72 | 84.06 |

### Accuracy and robustness of Noisy Turnpike solutions on synthetic instances

We tested how accurately the MM (Section 2.2), backtracking, distribution matching [12], and gradient descent methods were able to reconstruct point sets. Table 2 shows the median MAE normalized by the uncertainty for a representative set of problem sizes and uncertainties. Each method had 1 hour to solve each instance, with the exception of 10,000 and 100,000 point instances, which were given 90 and 6000 minutes respectively. The backtracking method was able to solve instances with 1,000 or fewer points, but exhibited larger errors than our method. The gradient descent method solved all instances with 500 or fewer points with residual error that ranged between 10 and 1,000 times higher than the MM approach. The distribution matching method performed similarly to our method but could not scale past 100 points. Our method was able to solve instances with 2,000 points with a median MAE that is 10 times lower than the uncertainty level up to a magnitude of 10^−4^, after which the scaling becomes distribution dependent.

**Table 2:** Median MAE normalized by the magnitude of measurement uncertainty ϵ across different point set sizes and uncertainties. We compare our method (MM), distribution matching (DM), backtracking (BT), and gradient descent (GD) approaches. A dash indicates that a method did not finish solving any instances of this size due to either memory or runtime constraints.

| $n \times 10^2$ | $10^{-6}$ | | | | $10^{-5}$ | | | | $10^{-4}$ | | | |
| --- | --- | --- | --- | --- | --- | --- | --- | --- | --- | --- | --- | --- |
|  | MM | DM | BT | GD | MM | DM | BT | GD | MM | DM | BT | GD |
| 1 | 0.2 | 0.49 | 26.4 | 1270 | 0.19 | 0.51 | 26.6 | 186 | 2.09 | 0.68 | 56.7 | 1.54 |
| 2 | 0.1 | — | 26.4 | 462 | 0.14 | — | 26.5 | 39 | 1.54 | — | 55.7 | 5.72 |
| 5 | 0.11 | — | 36.4 | 312 | 0.09 | — | 31.5 | 40 | 0.94 | — | 34.7 | 3.08 |
| 10 | 0.06 | — | 36.4 | — | 0.04 | — | 36.5 | — | 0.08 | — | 36.7 | — |
| 50 | 0.08 | — | — | — | 0.08 | — | — | — | 0.86 | — | — | — |
| 100 | 0.08 | — | — | — | 0.11 | — | — | — | 0.82 | — | — | — |
| 1000 | 0.12 | — | — | — | 0.08 | — | — | — | 0.11 | — | — | — |

Fig. 2 shows our method’s MAE and MSE over all settings plotted with respect to the magnitude of uncertainty. We observed that uncertainty in the distances correlated with reconstruction error, but the MAE is an order of magnitude lower than the uncertainty on average when the uncertainty is 10^−4^ or less. Instance size also affects the method’s error scaling. As the sample size varied between 50 and 100,000 points, the median MAE shown in Table 2 demonstrates a downward trend for fixed error rates. This suggests the method scales at least as well as it does on small point sets as the number of points grows. This is because the distance measurement linear system is highly overdetermined, which makes it resilient to uncertainty [19]. Last, we report the mean and standard deviation of the runtimes of different solvers in Table 4. The MM mean runtime was lowest across all point sizes and MM is the only method that successfully solved the 5,000, 10,000, and 100,000 point instances.

**Fig. 2:**
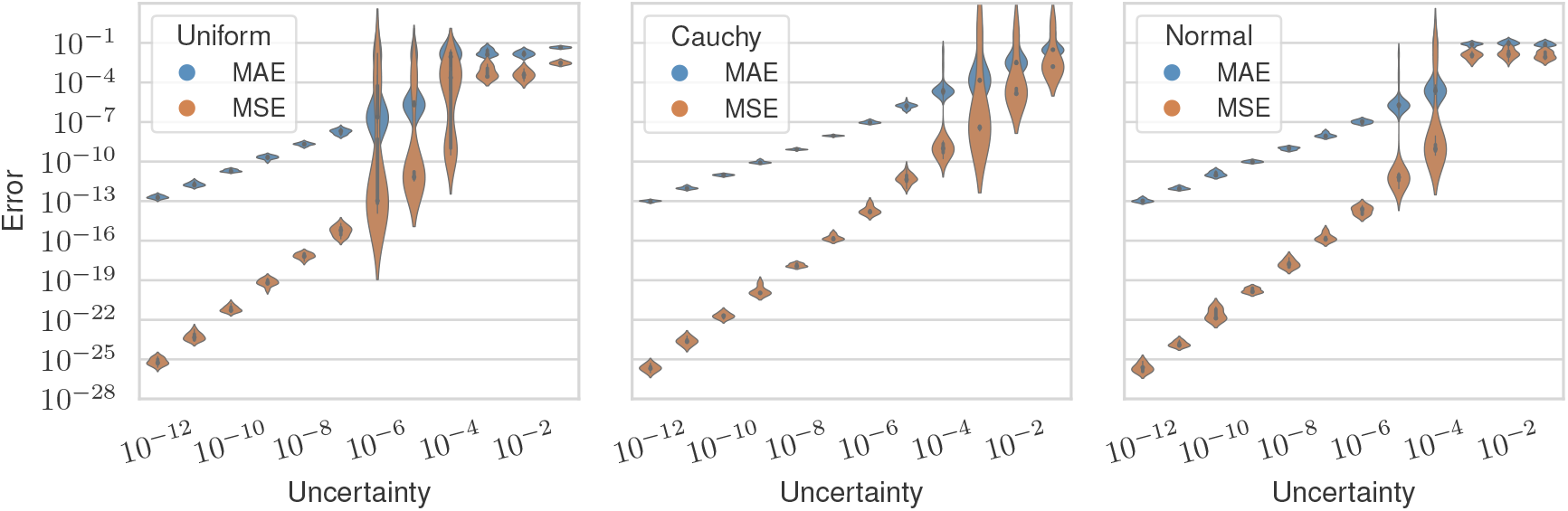
Mean absolute error (blue) and mean squared error (orange) between the estimated and ground vectors across all levels of measurement uncertainty and separated by distribution.

### Partial digestion experiments

We test the use of our algorithms for reconstructing genomes in the setting that uses an enzyme to partially digest DNA fragments at restriction sites [2]. The fragment lengths give the distances between all restriction sites, which are at unknown positions. The genome is assembled from the fragments after inferring the restriction site locations from the distances, a process equivalent to solving the Turnpike problem for linear genomes and the Beltway problem for circular genomes [12]. We simulated partial digestion instances to test our algorithms. For Turnpike instances, we used the human X chromosome’s centromere, and for Beltway instances, we used the full genome of the bacteria *Carsonella ruddii*. In both cases, we used 15-base-long enzymes and simulated the digestion process by sampling the DNA sequence such that each restriction site occurred between 10 and 500 times. We obtained digested DNA fragments by splitting the sequence at all of its occurrences. We added a signed Poisson random vector parameterized by a value λ > 0. This simulates when the enzymes cut too many or too few bases, both frequent occurrences in practice [6]. The Turnpike experiments were performed with our method and the gradient descent baseline, as they timed out for the other methods due to the size and amount of uncertainty. The Beltway experiments were performed with our algorithm and the distribution matching algorithm, as they are the only ones designed for uncertain Beltway instances.

Table 3 shows normalized MAE for the Turnpike experiments, which were performed with 10 to 2669 fragments. Our method recovered fragment locations with an MAE that scaled linearly to the uncertainty present in the measurements, performing orders of magnitude better than the gradient descent baseline. Table 5 shows normalized MAE for the Beltway experiments, which were performed with 10 to 54 fragments. Our method performed competitively with the distribution matching approach. In both cases, the generated point sets had highly repetitive distances, as the simulated restriction sites were frequently equidistant from one another, showing our method’s robustness to symmetric distances.

**Table 3:** Normalized MAE for 10 sizes and 4 uncertainty magnitudes using the MM and gradient descent (GD) algorithms on simulated partial digestions of a linear genome. A dash indicates that the algorithm did not finish due to memory constraints or runtime constraints.

| n | MM<br>($10^{-7}$) | GD<br>($10^{-7}$) | MM<br>($10^{-6}$) | GD<br>($10^{-6}$) | MM<br>($10^{-5}$) | GD<br>($10^{-5}$) | MM<br>($10^{-4}$) | GD<br>($10^{-4}$) |
| --- | --- | --- | --- | --- | --- | --- | --- | --- |
| 10 | 0.152 | $2.31 \times 10^6$ | 0.148 | $1.54 \times 10^5$ | 0.163 | $1.70 \times 10^4$ | 0.142 | $1.34 \times 10^3$ |
| 64 | 0.213 | $1.89 \times 10^5$ | 0.219 | $6.42 \times 10^4$ | 0.220 | $1.87 \times 10^3$ | 0.232 | $4.20 \times 10^2$ |
| 142 | 0.358 | $8.22 \times 10^5$ | 0.359 | $3.44 \times 10^4$ | 0.357 | $9.28 \times 10^3$ | 0.365 | $6.12 \times 10^2$ |
| 183 | 0.282 | $1.28 \times 10^6$ | 0.268 | $1.75 \times 10^5$ | 0.288 | $9.37 \times 10^3$ | 0.290 | $1.34 \times 10^3$ |
| 530 | 0.071 | — | 0.070 | — | 0.080 | — | 0.066 | — |
| 959 | 0.119 | — | 0.121 | — | 0.125 | — | 0.139 | — |
| 1209 | 0.119 | — | 0.132 | — | 0.127 | — | 0.115 | — |
| 1451 | 0.059 | — | 0.061 | — | 0.043 | — | 0.072 | — |
| 2669 | 0.048 | — | 0.039 | — | 0.050 | — | 0.048 | — |

**Table 4:**
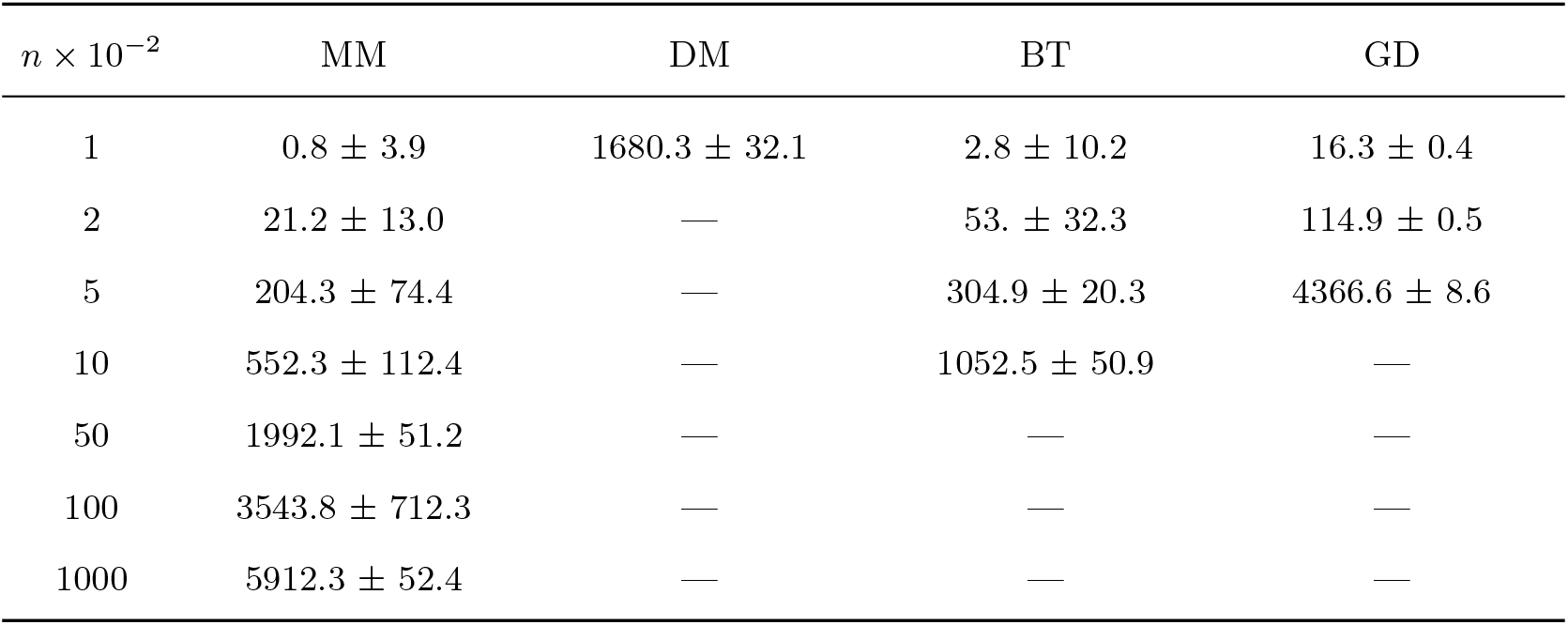
Mean runtime in seconds with standard deviations across point sizes for various Turnpike solvers.

**Table 5:**
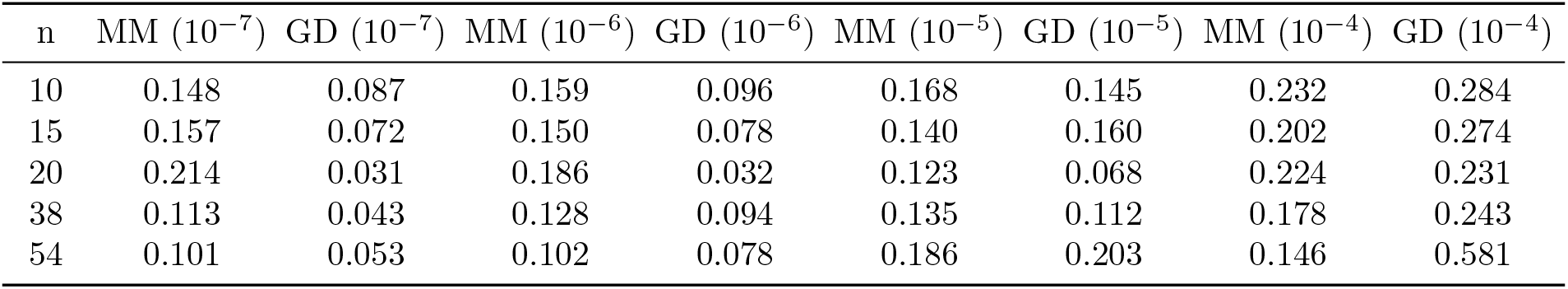
Normalized MAE for 5 sizes and 4 uncertainty magnitudes using the MM and distribution matching (DM) algorithms on simulated partial digestions of a circular genome.

### Labeled Partial Digestion Experiment

Pandurangan et al. [20] performed a labeled partial digestion problem (LPDP) recovery experiment using the restriction sites of the enzyme HindIII on the bacteriophage λ. For each distance *d*, they simulated relative uncertainty of order *r* ∈ [0, 1] by replacing *d* with a uniformly sampled integer in [(1 − *r*) *d*, (1 + *r*) *d*]. They varied *r* between 0% and 5% to mimic experimental settings, where 2% to 5% is expected. Each experiment was repeated 100 times and was reported as a success when the recovered distances were within the relative uncertainty of the ground truth set. We repeated this experiment using our base algorithm (MM; without using additional labeling information) and our partition-update formulation given in Alg. 5 of Appendix C. The results are shown in Table 6, where we see our method performs competitively without additional labels and further improves when provided with the labels. All instances ran in less than one second across all uncertainties and solvers.

**Table 6:** Recovery success for % relative error for our base sovler (MM), our partition solver (PMM), and Pandurangan et al.’s solver [20].

| $r$ | MM | PMM | LPDP |
| --- | --- | --- | --- |
| 0% | 100% | 100% | 100% |
| 1% | 99% | 99% | 98% |
| 2% | 96% | 97% | 96% |
| 3% | 95% | 96% | 94% |
| 4% | 92% | 94% | 91% |
| 5% | 89% | 92% | 87% |

## 4 Conclusion

Noisy Beltway and Noisy Turnpike are NP-hard problems that aim to recover a set of onedimensional points based on a corrupted pairwise distances. These problems find application in widespread biological contexts. We introduced a novel optimization formulation and an alternating algorithm built from sorting and implicit matrix multiplication. This leads to an asymptotic runtime of *𝒪*(*n*^2^ log n) time per iteration with *𝒪*(n) auxiliary memory. To escape low-quality local optima, we introduced a divide-and-conquer step to fix common errors. We performed large-scale experiments with approximately 25 billion distances (equivalent to 100,000 points) to showcase the efficiency of our method. In contrast, previous methods are infeasible with as few as 125,000 distances (equivalent to 500 points). We also demonstrated the method’s robustness and scalability in a variety of challenging situations, including large-scale uncertainty and distance duplication. Our algorithm efficiently solves large distance sets with realistic levels of uncertainty, opening up new avenues of research into biological applications of Turnpike and computational geometry problems.

## Acknowledgements

This work was supported in part by the US National Science Foundation [DBI-1937540, III-2232121], the US National Institutes of Health [R01HG012470] and by the generosity of Eric and Wendy Schmidt by recommendation of the Schmidt Futures program. Conflict of Interest: C.K. is a co-founder of Ocean Genomics, Inc.

## A Matrix-free oracles to avoid storage cost

### Algorithm 4

*Q* and *Q*^⊤^ multiplication oracles

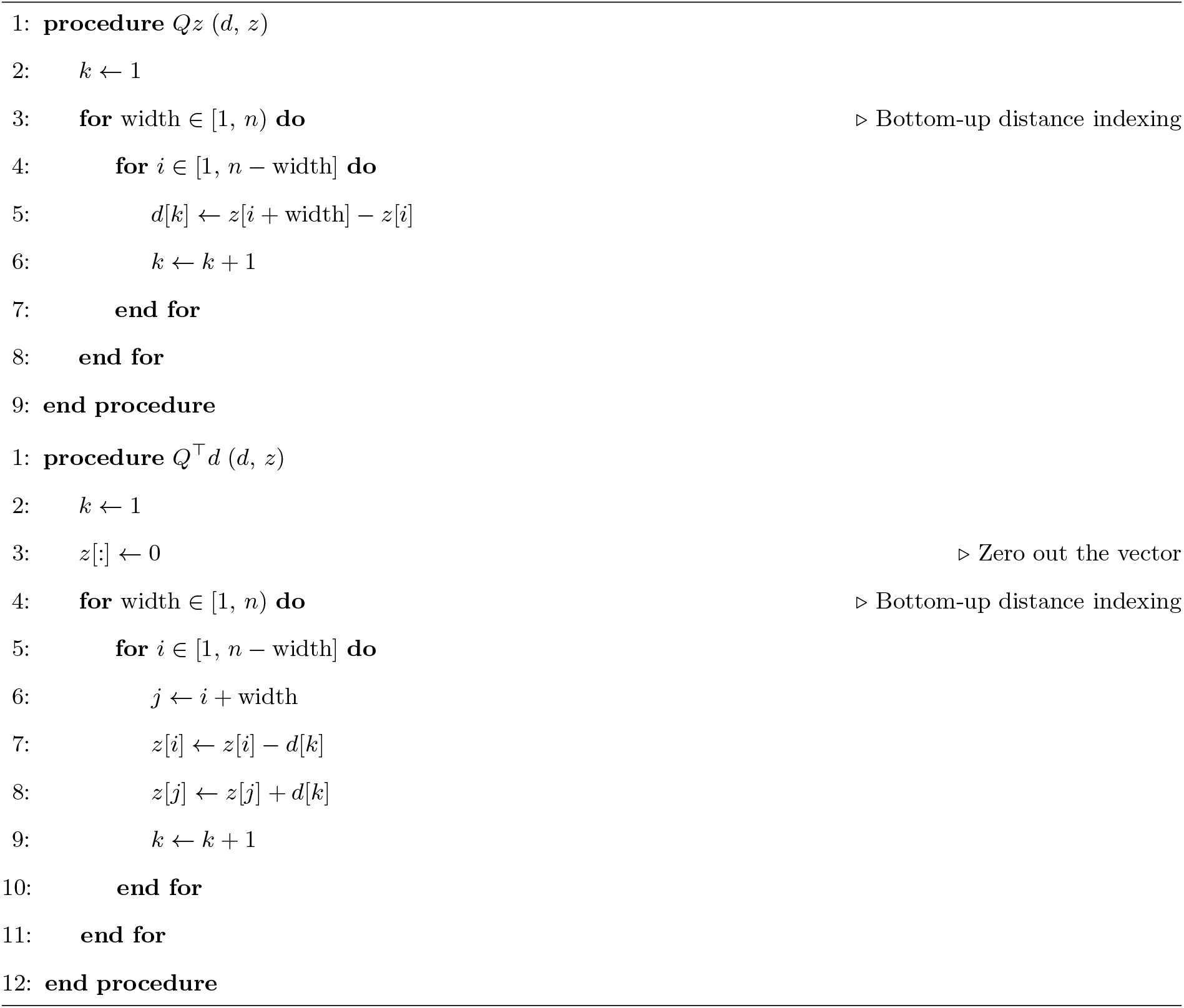

## B Visualizing behavior of the divide-and-conquer heuristic

**Fig. 3:**
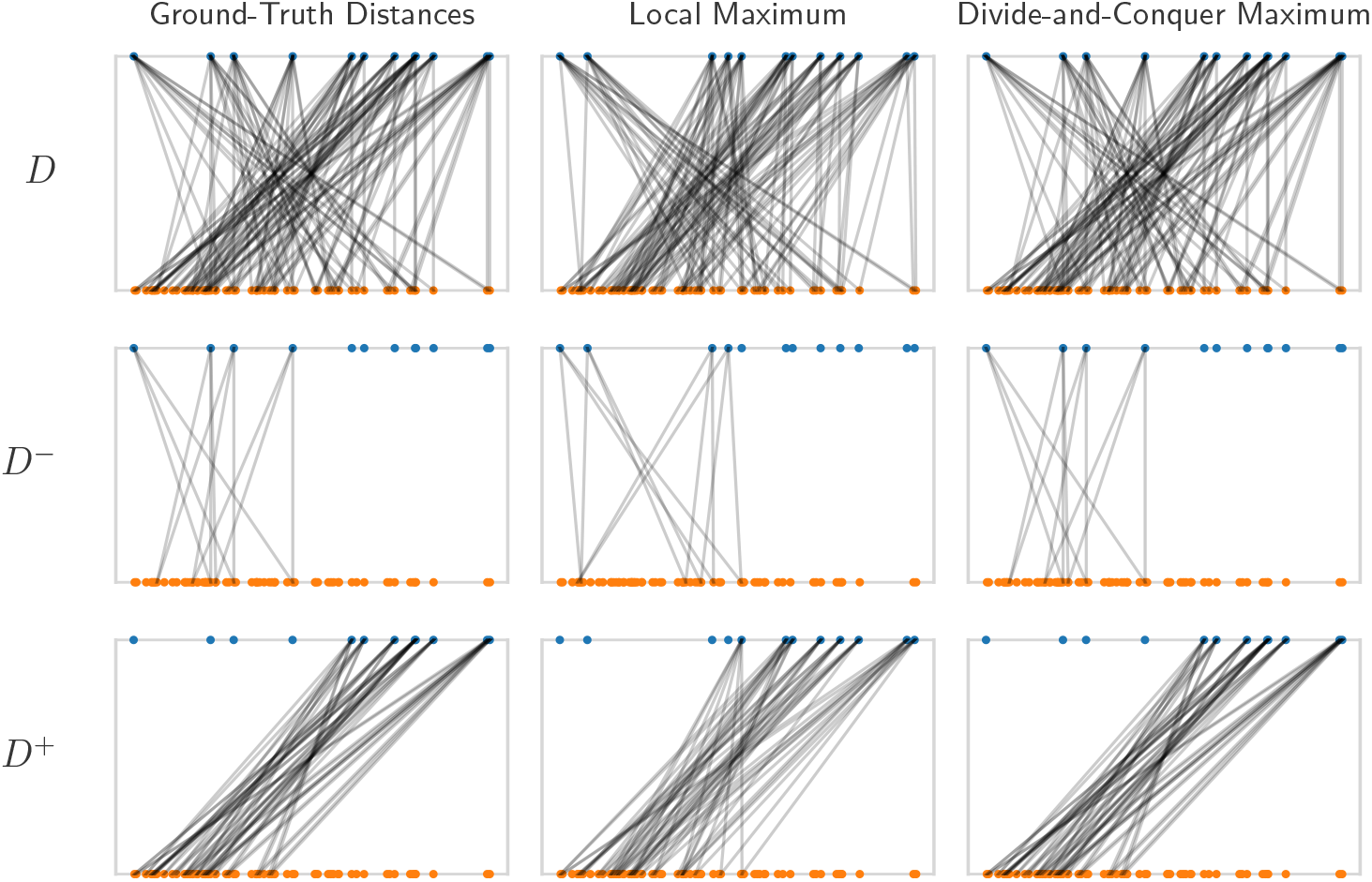
An example of the divide-and-conquer strategy applied to a local maximum, leading to global optimality. The figure presents a series of plots, where the point vector (blue) at the top corresponds to the distance vector (orange) at the bottom. Each distance has two edges to the points that generated it. From top to bottom, the rows represent the full distance set, the distance set associated with negative points, and the distance set associated with non-negative points. From left to right, we see the ground truth matching, local maximum matching, and divide-and-conquer maximum matching. This instance displays the ideal case for the divide-and-conquer approach, where each half of the interval gets nearly its entire distance set.

## C Extension to Problem Variants

### C.1 Beltway

We cast the Beltway problem as solving for the angle vector 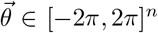 that contains the angles between every two points. This has precisely the same symmetry-breaking constraints as a linear point vector. We constrain the points to this set in order to cover the circle twice using two different orientations. With this setup, the existence of any solution guarantees that there exists at least one solution with an angle vector 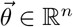 where *θ*_1_ = 0 because a rigid rotation does not change the pairwise distances. Similar to the Turnpike case, we can also freely reorder the angles without changing the problem, so we constrain our search to a sorted angle vector *θ*_1_ ≤ *θ*_2_ ≤ … ≤ *θ*_*n*_.

We account for the two different arcs between any two points by solving for a 2*n* dimensional angle vector indexed by integers in [−*n, n*]−{0}. Let this new vector be defined as the concatenation 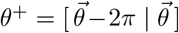, which constrains our search by the relationship between the clockwise and counter-clockwise arcs. This fixes the ambiguity problem, as both arclength distances are in the standard linear distance set of this vector. To see this, we have for *i* < *j* that 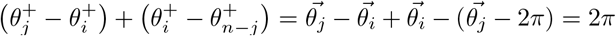, i.e., the entire partition of the circle is present. We also maintain sorted order because 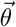 is sorted, 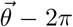 is sorted, and 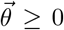. Every distance is duplicated because every point has two representatives, so we have to duplicate the distance set before running the algorithm. There are also *n* distances of the form *θ*_*k*_ − *θ*_−*n*+*k*_ missing, but these are simply the circumference of the circle, i.e., 2π. Alg. 2 can be modified to correctly assign these extra distances with a single conditional.

### C.2 General Partial Digest Problems

The labeled partial digest problem (LPDP) [20] is a variant of the Turnpike problem where we receive both the endpoint distances 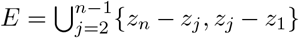 and the all-pairs distance set *D*. The simplified partial digest problem (SPDP) [4] provides an adjacent distance set 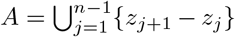 and the same endpoint distance set *E*. For LPDP, can use the base procedure of Algorithm 1, but that does not make efficient use of all available information. For SPDP, we can use a construction similar to the Beltway formulation, but that also does not efficiently use the additional labeling information. To that end, we abstract our method to work with any possible one-dimensional distance assignment problem via the unifying method of distance set partitions. To use this framework, the input to every problem must be turned into a partition. In the Noisy case, optimal assignments with respect to a structured convex objective function like the *𝓁*_1_ norm can be found using fast *𝒪* (*n* log *n*) projection algorithms [15]. For example, LPDP requires us to form *D* − *E* from *D* and *E*, and SPDP requires us to form the partition *A* − *E, E* − *A*, and *A* ∩ *E*. Additionally, general one-dimensional distance assignments can require the simulation of missing distances, e.g. *D* − (*A* ∪ *E*) for SPDP. Our framework accommodates for this by filling in these distances using the estimated point vector from the previous iteration, a process we describe below.

#### Partition formulation

Suppose we receive as input a partitioned distance set *D* = *D*^1^ ⊔ … ⊔ *D*^*s*^ and a partition of the linear indices {1, …, *m*} = *I*^1^ ⊔ … ⊔ *I*^*s*^ such that the submatrix 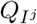 contains the measurements that generated the distance subset *D*^*j*^. (We use − to indicate a union of two disjoint sets.) For each *D*^*j*^, we will optimize for a permutation matrix *P* ^*j*^ of size |*I*^*j*^| × |*I*^*j*^| that rearranges *D*^*j*^. To do this, we replace the assumption that the distance vector *D* is sorted and instead assume each component of the partition *D*^*j*^ is sorted. With these changes, our new objective becomes

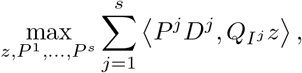

where *z* is a centered, sorted point vector. Suppose we fix all permutation matrices except for *P*^*j*^. All terms not involving *P*^*j*^ are constant in this subproblem, so we drop them without change in optimality. With this, the objective simplifies to

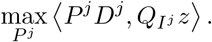

From Lemma 1 and Proposition 1, we know an optimal choice of *P*^*j*^ is the transpose of a permutation matrix that puts 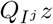 into sorted order. On the other hand, when the permutation matrices are fixed and *z* is not, the objective reduces to the original:

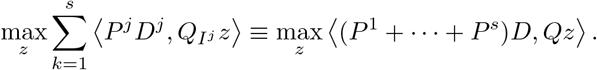

Note that we slightly abused notation by adding the permutations, but this is fine as we can simply lift each permutation to a partial permutation with zeros on all disjoint indices. Thus, an optimal choice is given by the same closed-form presented in Algorithm 1, i.e., *z* = unit [*Q*^⊤^(*P*^1^+ … + *P*^*s*^)*D*].

#### Missing distances

Suppose a subset of distances *D*^0^ are missing from the input. Letting *I*^0^ denote the indices of the missing subset, the index partition is then {1, …, *m*} = *I*^0^ ⊔ *I*^1^ ⊔ … ⊔ *I*^*s*^. Our formulation can accommodate for these missing entries with a simple change: we use the current estimate *z*^*t*^ to simulate them. Note we only need these distances for estimating *z*^*t*+1^, as the matching subproblem using these distances results in no change.

#### Block-Update Algorithm

We have a closed-form solution to every subproblem, and the objective is non-decreasing with respect to each closed subproblem. Thus, we can use block updates (i.e., update one subproblem at a time) or parallel updates (i.e., update and aggregate all subproblems) to make progress. For simplicity, the version we present (Algorithm 5 below) is an extension of Algorithm 1 that updates all permutations in parallel and then updates the point vector. As before, we stop the optimization loop once the vector reaches a fixed-point. We also adapt Algorithm 2 to run on the partitioned distance set while preserving the *𝒪* (*n*) space complexity, provided the partition index of an interval can be queried efficiently (described below).

##### Algorithm 5

Block-Descent Minorization-Maximization (MM)

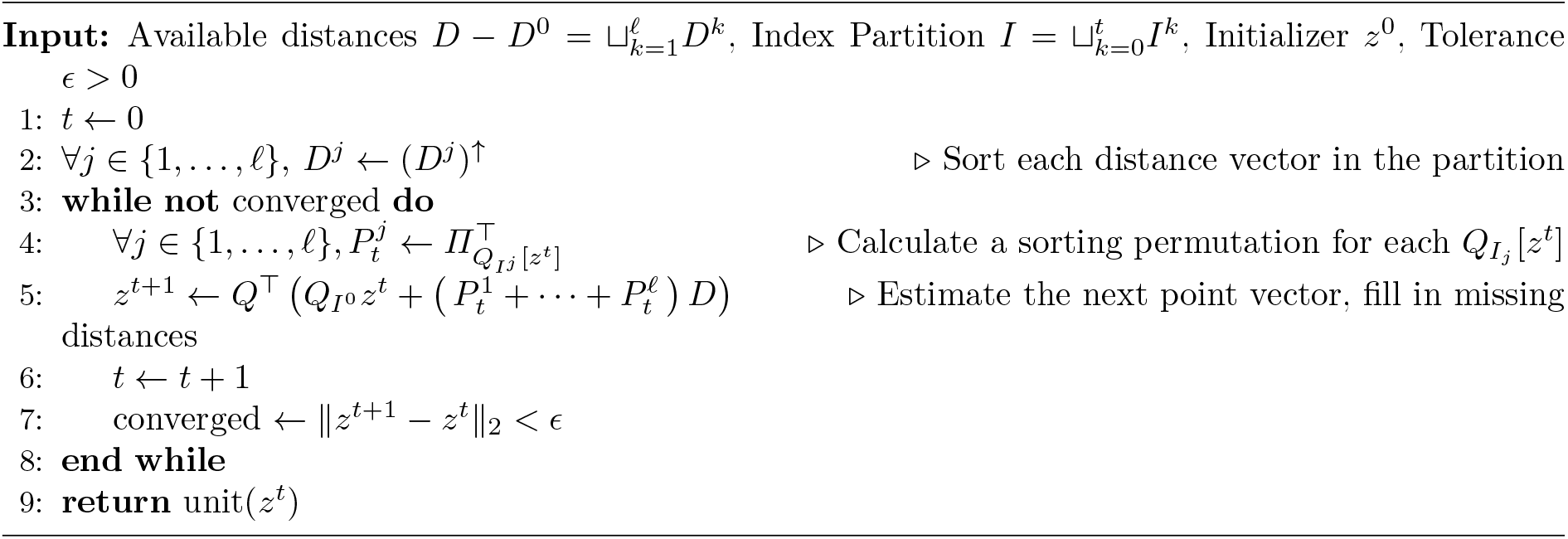

##### Algorithm 6

*Q*^⊤^ applied to matched *D* with a partition

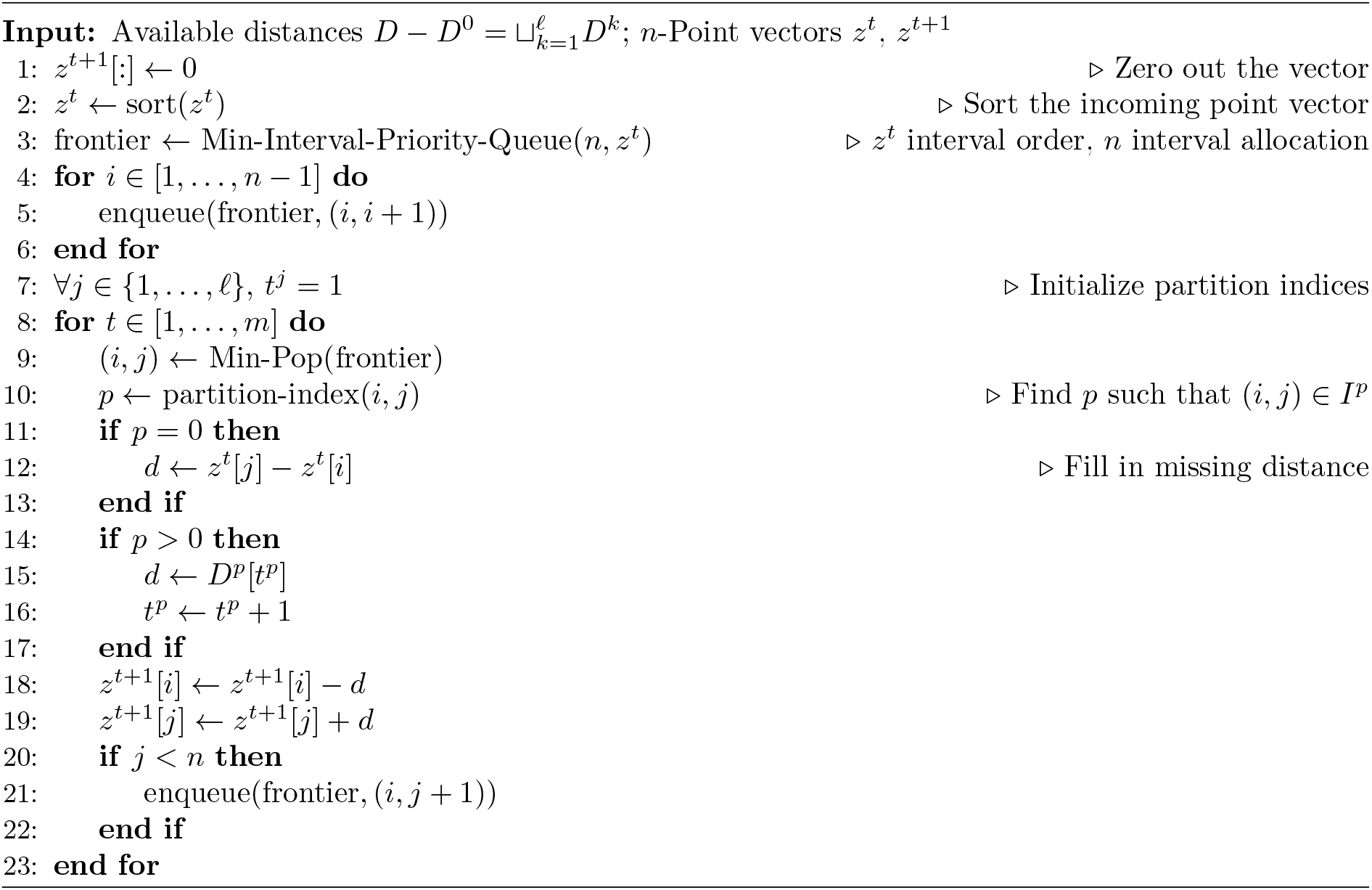

There are only three differences between Algorithm 3 and Algorithm 6.

1. Before the loop, we initialize indices *t*^1^, …, *t*^ℓ^ for the sets in the available partition.
2. When an interval with a missing distance is relaxed, we simulate the distance from *z*^*t*^.
3. When an interval in *I*^*j*^ with *j*≠0 is relaxed, we use the distance *D*^*j*^[*t*^*j*^] and set *t*^*j*^ ← *t*^*j*^ + 1.

These changes ensure a distance from the correct partition segment is assigned to the current interval, and if it is missing, it is filled in from the current vector *z*^*t*^. These changes do not affect the asymptotic runtime except for potentially checking an interval’s partition index. This step is run once per iteration, so the runtime becomes *𝒪* (*n*^2^ log *n* + *pn*^2^), where *p* is the cost of checking the partition index. For natural partitions, checking the index takes constant time and requires no additional space, leaving both the runtime and space complexity unchanged. For example, in LPDP, we only check whether the current interval has endpoint 1 or *n*, and in SPDP, we check if the interval is from two adjacent points, if it is from an endpoint distance, or if it is from neither.

